# Domain-Invariant Feature Learning for Patient-Level Phenotype Prediction from Single-Cell Data

**DOI:** 10.1101/2025.09.22.677881

**Authors:** Mathias Perez, Justin Hong, Aaron Zweig, Elham Azizi

**Author notes:** These authors contributed equally (Mathias Perez, Justin Hong). {, }.

## Abstract

Accurate prediction of patient-level disease status from single-cell RNA sequencing (scRNA-seq) data is critical to enabling precision diagnostics. However, study-specific artifacts induce spurious correlations that limit generalization and interpretability. We studied this problem in the context of Multiple Instance Learning (MIL), a framework where each patient is modeled as a set of single-cell profiles. To improve robustness to domain shifts, we propose an adversarial and metric-based approach that learns domain-invariant representations while preserving task-relevant biological variation. We benchmarked our method on a systemic lupus erythematosus (SLE) dataset with synthetically added spurious features and evaluated its performance on two real-world scRNA-seq atlases: a cross-tissue immune dataset and a COVID-19 severity atlas. Across all settings, we observed consistent improvements in out-of-domain accuracy and more biologically faithful model attributions. Our findings establish a new standard for robust, interpretable patient-level prediction under domain shifts using scRNA-seq.

## 1 Introduction

Single-cell RNA sequencing (scRNA-seq) enables high-resolution profiling of gene expression at the cellular level and has become a cornerstone in modern biomedical research. While most applications have focused on cell-level analyses, an emerging and highly promising direction is patient-level disease classification [1, 2, 3, 4], where each patient is modeled as an unordered set of cells. This formulation opens the door to building interpretable, generalizable models that could not only improve diagnostic performance but also offer deeper insights into disease mechanisms. However, like many tasks involving multi-study data integration, patient-level prediction faces a key challenge: spurious correlations introduced by batch effects, demographic shifts, or protocol differences. These correlations can lead models to rely on non-causal features, resulting in poor generalization and misleading interpretations. To address this, we propose a domain-adversarial strategy to enforce representation invariance across environments (**Figure 1**). Our contributions are threefold: (1) we systematically evaluated the limitations of existing domain generalization methods under controlled spurious correlations; (2) we introduced a bag-level, Conditional Domain-Adversarial Neural Network (CDANN)-based approach enhanced with CenterLoss to promote intra-class compactness [5, 6]; and (3) we demonstrated improved out-of-domain accuracy and more stable feature attributions on both a semi-synthetic benchmark and two public datasets.

**Figure 1.**
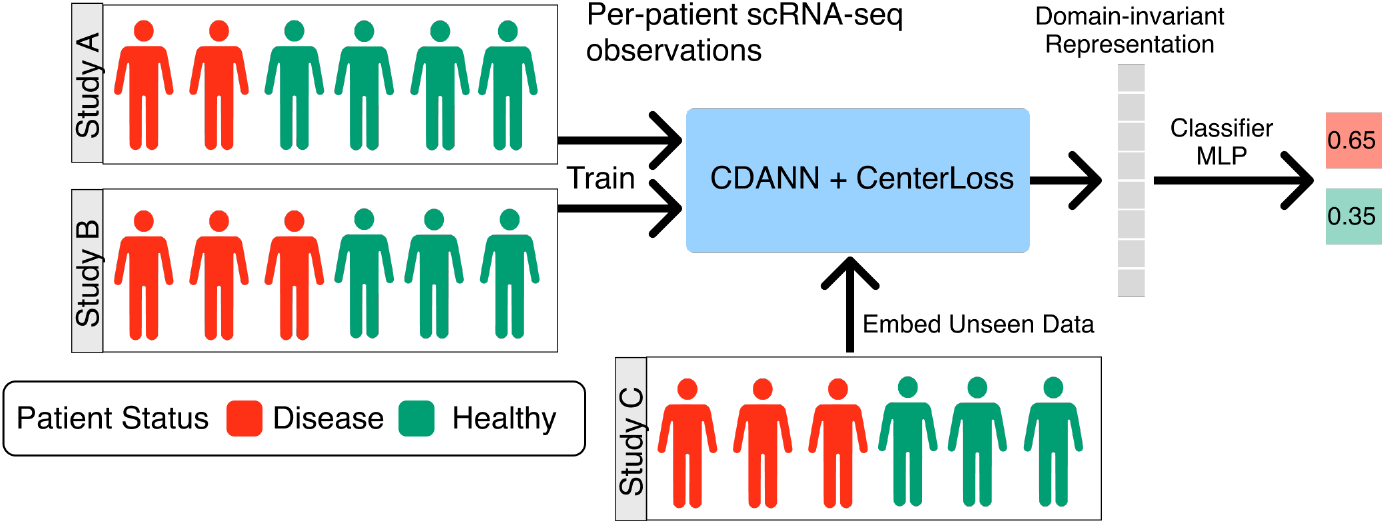
Illustration of problem setting and application of the proposed CDANN+CenterLoss approach. Provided a multi-study multi-patient scRNA-seq dataset, we aim to produce a domain-invariant representation of each patient for robust patient phenotype classification. By training over multiple studies/environments with plausible spurious features, we incentivize the model to learn relevant features shared across the environments.

## 2 Related Work

### 2.1 Multiple Instance Learning

Multiple Instance Learning (MIL) is a weakly supervised framework where labels are provided at the group (or “bag”) level, while individual instances remain unlabeled. Bags are treated as samples drawn from a latent distribution, and the outcome depends on a weighted combination of instance-level contributions. This perspective is well-suited to biological settings where subtle but informative signals are distributed across specific subpopulations of cells, e.g., transcriptional shifts in key cell types correspond to patient-level disease phenotypes.

In recent years, MIL has seen a shift toward embedding-level approaches, where each instance is first encoded into a vector and then aggregated into a bag-level representation via a permutation-invariant function. Foundational work like DeepSets [7] established theoretical guarantees for such models, proving that any permutation-invariant function can be decomposed as a sum over transformed instances. Building on this, attention-based pooling methods like AttMIL [8] introduced instance-level importance weighting, improving both flexibility and interpretability.

Further advances include the Set Transformer [9], which uses attention mechanisms to model higher-order interactions between instances, and SetNorm [10], which improves training stability and representation quality for high-dimensional data. These innovations have been widely adopted in computational biology, where the order of cells in a sample is irrelevant but their contextual relationships are crucial.

In our work, we adopted this modern embedding-based MIL formulation for patient-level disease classification from scRNA-seq data. We used DeepSets++ [10] as the core architecture for encoding sets of single-cell profiles into compact, informative patient-level embeddings.

### 2.2 Patient-level Phenotype Prediction

Patient-level modeling from scRNA-seq has been explored via MIL architectures such as CloudPred[1], ProtoCell4P[2], scMILD[3], and SingleDeep[4]. MIL is especially well-suited to this setting because each patient can be represented as a bag of cells, and the phenotype is determined by complex patterns across heterogeneous subpopulations rather than individual cells. By treating cells as instances and patients as bags, MIL enables direct patient-level prediction while also offering interpretable insights into which cellular subsets drive phenotypic variation. Existing approaches emphasize set-based aggregation and interpretability, but they are primarily based on empirical risk minimization (ERM) and typically do not address distributional shifts across cohorts or study sites.

Our work extends these approaches by introducing explicit domain generalization techniques into the MIL pipeline.

### 2.3 Invariant Learning

Learning invariant representations is central to domain generalization, especially in biological settings where technical effects, donor variability, or cohort shifts introduce spurious signals. A variety of methods have been developed to enforce robustness across environments, including risk-based regularization, adversarial alignment, and metric-based constraints, which we will now describe in more detail.

#### Risk-based Domain Generalization

Several recent methods attempt to improve robustness across training environments by modifying the training objective itself. These include Empirical Risk Minimization (ERM), Group Distributionally Robust Optimization (GroupDRO)[11], Invariant Risk Minimization (IRM)[12], and Risk Extrapolation (REx)[13]. All these methods operate by computing environment-specific risks and adjusting the optimization objective to promote generalization.

#### Adversarial Domain Alignment

While classic domain generalization methods such as IRM, GroupDRO, and REx aim to encourage invariance to spurious correlations, recent analyses suggest they may fall short in fully removing them from the learned representations. In particular, [14] shows that these methods often highlight core features no better than ERM, and primarily act by reweighting, rather than removing spurious signals. This is problematic in high-stakes applications such as biomedical prediction, where reliance on non-causal features can undermine generalization and interpretability. To address this, adversarial strategies seek to explicitly erase environment-specific information from the representation.

Adversarial domain adaptation methods such as Domain-Adversarial Neural Networks (DANN) [15] introduce a domain classifier *d* trained to predict the source environment from the learned features *ϕ*(*x*), while the feature encoder *ϕ* is trained adversarially to fool *d* via a gradient reversal layer:

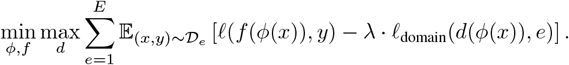

This results in domain-invariant embeddings. However, aligning all domains can erase task-relevant differences (e.g., disease signal in biology).

To remedy this, Conditional DANN (CDANN) [5] conditions the domain classifier on the predicted label, preserving class semantics during alignment. CDANN is especially useful in settings with label-dependent domain shifts, such as cell-type composition differences across patients. In our setting, we apply the conditioning to patient-level predictions.

#### Metric Learning Losses

Beyond domain alignment, robust feature learning can be enhanced by metric-based constraints that shape the geometry of the embedding space. Two notable examples are ArcFace[16] and CenterLoss[6], which encourage angular separation between classes and intra-class compactness, respectively. While ArcFace introduces an angular margin between class embeddings, CenterLoss penalizes deviation from class-specific centroids. The full formulations of these objectives are presented in Section 4.

In this work, we combined CDANN with CenterLoss to jointly promote domain invariance and class discriminability, aiming for robust, interpretable patient-level prediction in scRNA-seq data.

## 3 Methods

### 3.1 Notation and Model Components

We model each patient indexed by *i* as a *bag of n*_*i*_ *cells* with cells indexed by *j*, represented as a multiset:

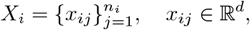

where *n*_*i*_ is the total bag size, with a corresponding disease label *y*_*i*_ ∈ [1 … *K*]. Each bag is sampled from a domain (or environment) *e* ∈ ℰ, such that the training dataset is partitioned into *N*_*e*_ domain-specific subsets:

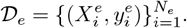

We aim to minimize the *test domain risk*:

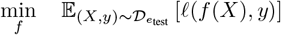

for a learned model *f* and classification loss *ℓ*, while having access only to data from a subset of domains ℰ _train_ during training. This setup falls under the *domain generalization* setting, where the test environment is unknown a priori.

### 3.2 CDANN with CenterLoss

Our model combines Conditional Domain-Adversarial Neural Networks (CDANN) with CenterLoss to promote domain invariance and class compactness. Given a bag *X*_*i*_, the encoder *ϕ* produces an embedding *z*_*i*_ = *ϕ*(*X*_*i*_). We use the permutation-invariant DeepSets++ [10] model class for *ϕ*. The loss comprises three terms:

#### Classification Loss

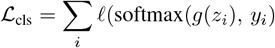

where *g*(·) is a small multi-layer perceptron (MLP) mapping embeddings *z*_*i*_ to a logit vector. The cross-entropy loss *ℓ* then encourages the predicted class to match the true disease label *y*_*i*_.

#### Adversarial Domain Loss

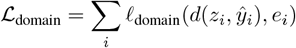

The domain discriminator *d* is conditioned on the predicted label *ŷ* _*i*_, forcing the encoder to remove environment-specific information.

#### Center Loss

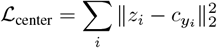

This term encourages embeddings *z*_*i*_ to cluster around their respective class centers 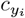. Each 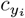 denotes the learnable vector associated with class *y*_*i*_. In practice, the centers are updated after each batch by moving them towards the empirical mean of their assigned embeddings with a momentum-like rule:

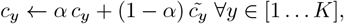

where 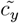 is the class centroid over the batch and *α* ∈ [0, 1] controls the update rate. The total training objective is:

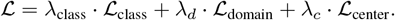

We explored different scheduling strategies for the coefficients *λ* _*d*_ and *λ* _class_ to improve training stability. In particular, we warm-started the model by initially disabling the adversarial loss (i.e., *λ*_*d*_ = 0), allowing the classifier to first learn task-relevant features. After this warm-up phase, we gradually increased *λ*_*d*_ while alternating updates between the classifier and the domain discriminator.

This procedure avoided the common collapse scenario where early adversarial updates overpowered a still-untrained classifier.

## 4 Results

### 4.1 Setup and Problem Formulation

In real-world biological datasets, domain shifts arise naturally due to demographic variation, technical effects, or study-specific factors. These shifts may cause *spurious correlations* between non-causal features and the prediction target. While a model trained on multiple environments might learn to ignore such spurious signals if all environments are seen during training, its performance can degrade dramatically when evaluated on a held-out domain where spurious correlations differ.

To systematically study this challenge, we built two controlled settings where certain environments contain features that are strongly or weakly correlated with the label, but these associations are absent or inverted in others. We created four environments as follows: (1), (2), and (3) are used for training/validation where the spurious feature is either strongly or weakly correlated with the label, while environment (4) contains an *anti-correlated spurious feature* and serves as the test domain (**Supp. Figure 1**). This setup highlights the failure modes of models that rely too heavily on features that are not consistent predictors across training environments.

For real datasets, we did not have access to ground-truth spurious features. Instead, we constructed environments based on prior understanding of which patient-level metadata would be most likely to exhibit distribution shifts across groups.

### 4.2 Colored MNIST

Before applying our method to biological data, we first validated our approach on the well-known Colored MNIST benchmark. In this setting, the digit classification task was corrupted by a spurious feature—color—which is strongly correlated with the label during training but inverted at test time.

We reproduced this setup by assigning each digit a specific color in the training environments (e.g., red for digit 1, green for digit 7) and flipping the color-label association in the test environment. This simulated a domain shift where the spurious cue becomes misleading. Our goal was to test whether metric learning losses and adversarial training could prevent reliance on this spurious feature.

Standard ERM (**Appendix 6.2**) achieved high training accuracy but completely failed under spurious inversion, with test accuracy dropping to 7% (see **Supp. Table 1**). In contrast, our CDANN + CenterLoss model restored performance to nearly 100% on the held-out domain. The UMAP visualizations (**Figure 2**) confirmed this: while ERM embeddings clustered by color, revealing dependence on the spurious feature, CDANN successfully removed this confounding, yielding clusters that aligned purely with digit identity.

**Table 1:** Test accuracy results across different datasets and methods. The best performing method is bolded for each dataset.

| Dataset | CDANN + CenterLoss | ERM | GroupDRO | REx |
| --- | --- | --- | --- | --- |
| Colored MNIST | **0.98** | 0.07 | 0.26 | 0.00 |
| Semi-synthetic SLE [17] (w/o spurious eff.) | **0.906** | 0.891 | 0.891 | **0.906** |
| Semi-synthetic SLE [17](w/ spurious eff.) | **0.634** | 0.00 | 0.008 | 0.052 |
| Cross-tissue Immune Atlas [18] | **0.795** | 0.682 | 0.773 | 0.773 |
| COVID-19 Atlas [19] | **0.712** | 0.654 | 0.552 | 0.677 |

**Figure 2.**
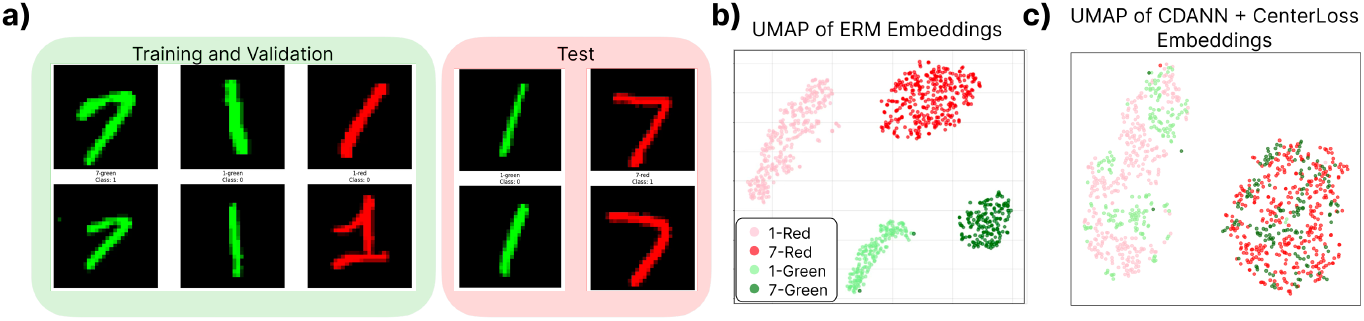
Colored MNIST proof-of-concept. (a) Dataset design with color as a spurious cue, flipped at test time. (b) UMAP of digit embeddings after ERM training: color dominates and induces clustered confounding. (c) UMAP after CDANN training: embeddings are class-discriminative, with no visible influence of color.

This experiment served as a clear validation of our approach: adversarial training combined with compactness-inducing regularization could effectively suppress spurious correlations and recover causal decision boundaries.

### 4.3 Semi-synthetic SLE benchmark

Next, we created a semi-synthetic variant of a Systemic Lupus Erythematosus (SLE) scRNA-seq dataset [17] (See **Appendix 6.3** for data preprocessing details). We defined environments based on patient age groups, leveraging the fact that disease severity in lupus is strongly age-dependent. This correlation introduced a natural spurious feature: any gene expression signal linked to age may confound disease prediction if not properly disentangled.

In selected source environments, we synthetically introduced a spurious correlation between the disease label and gene expression (**Supp. Figure 1**). Specifically, we defined class 1 as the positive class in our binary classification task (e.g., patients with the disease), and we injected a spurious signal by artificially modifying the expression level of a designated gene or gene set such that it became predictive of class 1 only within those environments. In the strong spurious setting, this was done by upregulating the gene(s) across all cells of class 1 patients in the spurious domains, while ensuring that this correlation is absent or even reversed in the held-out test domain. In the weak spurious setting, the same gene(s) were perturbed only in a subset of cells per patient, introducing within-patient heterogeneity and weakening the strength of the spurious association.

When no spurious correlations were introduced, all models, including ERM, GroupDRO, REx, and CDANN, achieved high predictive accuracy (**Table 1**) across all environments. This confirmed that the core task was learnable and that none of the models struggled when spurious features were either absent or consistent across domains. We initially included IRM in our experiments; however, consistent with prior reports, it exhibited severe optimization instabilities and failed to achieve meaningful accuracy. As a result, we do not report its results in Table 1.

In the presence of spurious correlations, particularly in the held-out domain where the spurious association was flipped, ERM and GroupDRO failed to generalize, with accuracy collapsing to nearly zero. REx showed some robustness but still performed poorly. In stark contrast, CDANN combined with CenterLoss remained robust and achieves over 60% accuracy in this challenging setting, without any tuning or data augmentation (**Table 1**).

These results highlighted that only CDANN combined with CenterLoss effectively ignored spurious signals and maintained high accuracy under distribution shifts. To better understand this robustness, we visualized the learned patient-level embeddings (**Figure 3**). Despite the presence of environment-specific confounding, CDANN learned a representation where patients clustered by class rather than by spurious environment, confirming its ability to learn invariant, task-relevant features.

**Figure 3.**
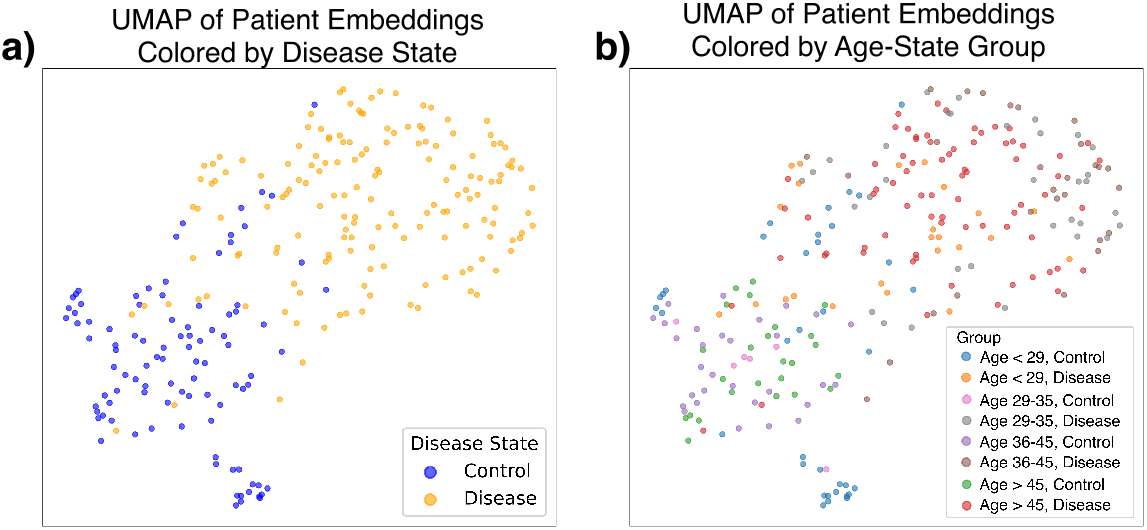
Patient-level embeddings learned by CDANN + Center Loss in the semi-synthetic SLE spurious setup. Color denotes class label. Patients clustered by class rather than by environment, indicating successful invariance to spurious domain-specific signals.

**Figure 4.**
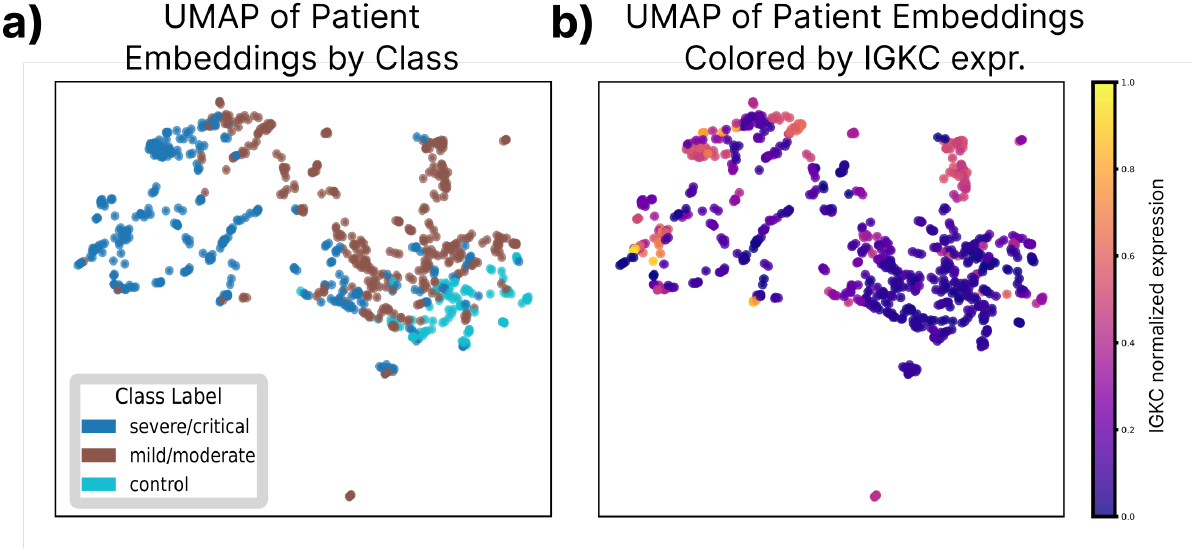
Gradient-based interpretation and gene expression signature of COVID-19 severity. **(a)** UMAP projection of patient samples colored by COVID-19 severity class. **(b)** The same UMAP projection colored by the normalized expression of *IGKC*. The expression *IGKC* (immunoglobulin *κ* constant region) tracks the severity gradient, consistent with plasmablast expansion in severe disease, suggesting its role as a molecular marker of control-to-moderate-to-severe transition.

Importantly, interpretability analysis revealed that once these regularization and adversarial components were added, the top contributing genes were more aligned with known SLE biology. For example, MHC Class II components (*HLA-DRB1, DQA1*) and FCGR3A were among the most influential features, consistent with autoimmune signaling pathways in SLE.

At the cell type level, ablation-based and DeepLIFT[20] methods (See **Appendix 6.5**) highlighted the role of plasmablasts and pDCs in disease prediction. Removing plasmablasts caused a sharp drop in disease classification probability, in line with their known role in autoantibody production. These results were more stable across training regimes compared to gradient-based attributions and DeepLIFT results.

### 4.4 Cross-tissue immune Atlas

The cross-tissue immune atlas [18] (see **Appendix 6.3** for data preprocessing details) profiles immune cells from 13 organs across 16 donors. Each bag corresponded to a donor–organ pair, and we defined the task to be predicting the organ of origin inferred from immune profiles. Donor-specific biases and organ heterogeneity introduced possible spurious associations. As such, patients were treated as environments and organs with 13-class labels. We subsampled each set of cells such that each bag contained ≤ 2,000 cells. Finally, we evaluated the models on held-out donor-organ pairs.

CDANN shrank the generalization gap from 29% to 17%—an absolute improvement of ∼ 11% on unseen donors (See detailed results in **Supp. Table 2**). Examining the UMAP embedding (**Figure 3**) showed that while ERM embeddings still clustered by *donor*, CDANN collapsed donor variation and revealed tight, organ-specific clusters.

### 4.5 COVID-19 Atlas

The COVID-19 atlas [19] (see **Appendix 6.3** for data preprocessing details) aggregates peripheral blood mononuclear cell (PBMC) profiles from 284 donors across six cities. Patients were grouped by collection site (city), and each bag represented the set of cells from one patient. Some cities exhibited site-specific biases (e.g., due to hospital protocols or demographic composition), allowing for implicit spurious correlations with the label, disease severity (*control, moderate, severe*). We held out one city for evaluation.

ERM again overfitted to site-specific batch effects, losing 27% from training to test (See **Supp. Table 3** for details. CDANN almost halved this regression and recovered 6% absolute accuracy, going up to 71% (See **Table 1**). Again, embedding plots illustrated the shift: ERM separated *cities*, whereas CDANN formed city-invariant severity gradients.

A DeepLIFT-based attribution (see **Supp. Figure 4**) identified *IGKC* (immunoglobulin *κ* light-chain constant region) as a key driver of the moderate-to-severe transition. Elevated *IGKC* reflects the expansion of plasmablasts/plasma cells. Consistently, *IGKC* is co-expressed with other immunoglobulin transcripts (e.g., *IGHG1, IGLC2, JCHAIN*) within plasmablasts/plasma cells [21], aligning with the surge of these cells in severe COVID-19 [22].

## 5 Discussion

To the best of our knowledge, this is the first work to introduce domain-adversarial learning for patient-level phenotype prediction using scRNA-seq data. Our results underscore the importance of domain generalization for patient-level disease prediction in scRNA-seq. Across both semi-synthetic and real-world datasets, we showed that standard methods such as ERM and GroupDRO can fail catastrophically in the presence of distribution shifts—particularly when spurious correlations invert across environments. By contrast, the CDANN model, augmented with CenterLoss, consistently improved generalization to held-out domains while yielding more biologically meaningful attributions.

### Robustness under controlled stress tests

Our semi-synthetic SLE benchmark revealed how easily classical methods can overfit to spurious structure. When artificial correlations were introduced and flipped at test time, both ERM and GroupDRO approached random performance, while CDANN+CenterLoss achieved over 60% accuracy. Importantly, these gains were obtained without any tuning to the held-out domain, validating the utility of domain-adversarial and metric-based regularization for causal feature learning.

### Interpretability and biological alignment

In all datasets, feature attributions produced by CDANN were more consistent with known biology. In the SLE dataset [17], top-ranked genes included MHC class II components and *FCGR3A*; in the COVID-19 atlas [19], our model identified an immunoglobulin gene (*IGKC*) and *CXCR2* as key severity markers, in line with recent literature on immune activation. These findings suggest that domain-invariant training not only improves accuracy but also enhances interpretability by steering models toward biologically grounded signals.

### Limitations and practical challenges

One key limitation of our evaluation lies in the artificiality of the semi-synthetic benchmark. While our setup provided a controlled environment to test generalization under label–spurious decoupling, the magnitude of spurious inversion may be stronger than typically observed in practice. Moreover, CDANN remains sensitive to hyperparameters and can collapse during training, particularly when class imbalance or noise is high. These instabilities pose a barrier to deployment in applied biomedical workflows.

### Future directions

Improving the stability and robustness of domain-invariant learning in weakly labeled, heterogeneous biological data remains an open challenge. Promising future directions include: (1) latent environment inference [23] to avoid reliance on explicit domain labels, (2) biologically informed priors [24] to guide regularization, and (3) hybrid contrastive-adversarial approaches [25] that combine robustness with representation richness. Additionally, incorporating self-supervised pretraining or cell-type–aware aggregation may improve model performance in low-data regimes.

In sum, our work provides a principled benchmark and methodological framework for tackling spurious correlations in patient-level scRNA-seq analysis. It highlights the limitations of conventional training pipelines and offers a concrete step toward robust and interpretable clinical prediction from high-dimensional single-cell data.

## Acknowledgements

We would like to thank Alessandro Grande and Ji Won Park for helpful feedback and discussions.

This work was supported by grant number 2022-253560 from the Chan Zuckerberg Initiative DAF, an advised fund of Silicon Valley Community Foundation.

## 6 Appendix

### 6.1 Additional comments on robust feature learning

Beyond domain alignment, robust feature learning can be enhanced by metric-based constraints that shape the geometry of the embedding space. Two notable examples are ArcFace [16] and CenterLoss [6].

ArcFace introduces an angular margin between class centers to promote discriminability:

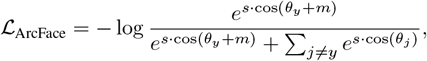

where *θ*_*y*_ is the angle between the input and its class center, *m* is a margin, and *s* is a scaling factor. This loss has been successful in face recognition but can be unstable under high class imbalance.

CenterLoss [6] directly penalizes the distance between embeddings and their class centroids:

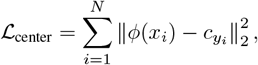

with 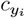 the learned center of class *y*_*i*_. This encourages intra-class compactness while complementing a standard classification loss. It is particularly useful for biological data where classes (e.g., disease severity) can be diffuse or overlapping.

### 6.2 Baselines and Training Objectives

To benchmark robustness, we compare four risk-based domain generalization methods. **Empirical Risk Minimization (ERM)** minimizes the average loss across environments,

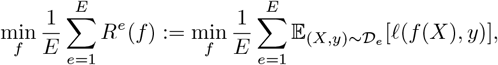

but is vulnerable to spurious correlations. **GroupDRO** mitigates this by optimizing for the worst-case environment risk,

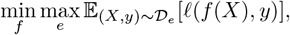

though it can be overly conservative. **Invariant Risk Minimization (IRM)** enforces invariance by penalizing the gradient of the loss with respect to a shared classifier across environments,

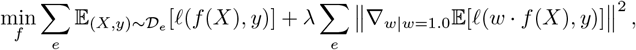

but suffers from optimization challenges. **Risk Extrapolation (REx)** promotes robustness by penalizing the variance of risks across environments:

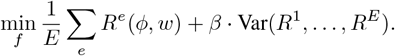

All baseline methods consistently failed to generalize under distribution shift on the semi-synthetic SLE dataset, performing poorly on held-out domains with inverted spurious correlations.

### 6.3 Data preprocessing

All datasets were obtained from the cellxgene portal under their respective publications [26]. To enable consistent input across cohorts, we restricted the feature space to the top 100 highly variable genes (HVGs) using the seurat_v3 flavor with the library UUID as the batch key.

For each dataset, raw count matrices were subset to the selected HVGs and stored as sparse matrices to reduce the memory footprint. Within each study (environment), counts were normalized on a per-cell basis using size-factor normalization to a fixed target library size (10^5^ counts per cell), followed by log1p transformation. This procedure ensured that technical effects such as sequencing depth were adjusted locally within each cohort.

### 6.4 Implementation details

All experiments were conducted using PyTorch with deterministic seeding and GPU acceleration. Models were trained at the patient (bag) level with a batch size of 32.

#### Model architecture

Unless otherwise specified, we used a DeepSets++ encoder with instance hidden dimension 32, followed by a bag-level MLP classifier with hidden size 64, dropout 0.4, and one hidden layer. Domain-adversarial models (DANN/CDANN) employed a discriminator with one hidden layer of size 32 and dropout 0.3. The number of output classes depended on the expected classification task. All linear layers were initialized using Xavier uniform initialization.

#### Training procedure

Models were trained for 200 epochs using Adam with a cyclical learning rate schedule (CyclicLR) between 10^−4^ and 10^−3^ (base 5 × 10^−4^), with a linear warm-up phase (`step_size_up` = 100). For CDANN, *λ*_class_ was kept at 1 during the entire training, while *λ*_adv_ was ramped with a logistic curve over the first 50 epochs with an added constant offset (+0.1) to stabilize early training. Entropy regularization was included with weight 1.0.

#### Regularizers

When CenterLoss was used, it was with weight *λ*_center_ = 1.0 and exponential update parameter *α* = 0.7. For IRM and REx, the variance/gradient penalties were weighted by *λ*_penalty_ = 0.5 and 1.0, respectively.

#### Warm-start and alternating strategies

In adversarial setups, we employed a warm-start strategy where the discriminator was initially unfrozen for 15 epochs, then alternated between frozen/unfrozen blocks of 35 and 15 epochs, respectively. This prevented the discriminator from overfitting and allowed the feature extractor to periodically refine representations without adversarial pressure.

Similarly, the class loss weight in CDANN was deliberately delayed to ensure that the shared feature extractor stabilized before being jointly optimized for class and domain discrimination.

### 6.5 DeepLIFT analysis

DeepLIFT (Learning Important Features through Propagating Activation Differences) [20] is a feature attribution method that decomposes the difference between a model’s output and that of a reference baseline into additive contributions from each input feature. In contrast to gradient-based saliency, which can be highly local and unstable, DeepLIFT propagates contribution scores through the network such that the sum of feature attributions matches the output difference relative to the baseline.

In our study, we applied DeepLIFT to two complementary settings: (i) gene-level attribution, where contributions are assigned to highly variable genes, and (ii) cell-type-level attribution, where contributions are aggregated across groups of cells belonging to the same type. Importantly, DeepLIFT provides *feature-wise* scores, that is, it identifies which input features (genes or aggregated cell-type profiles) are most influential for model predictions, rather than producing explanations tied to a specific patient.

The reference baseline was defined as the mean embedding across all cells in the dataset, ensuring that contributions were interpreted relative to an “average cell.” For each patient bag, DeepLIFT attributions were computed with respect to the true label, and absolute values of the scores were averaged across cells. This yielded (a) mean gene-level importance vectors for each class (control and diseased) as well as globally across patients, and (b) cell-type-level importance scores, obtained by averaging contributions over all cells of the same type. These aggregated results provided interpretable insights into which subsets of genes and cell types drive patient-level phenotype predictions.

## 6.6 Supplementary Tables

**Supplementary Table 1:** Test accuracy (%) of digit 7 with red color across different training scenarios. Columns correspond to color setups: (1) **Control**: only red digits (1 and 7) are seen during training, (2) **Red as spurious feature of digit 1**: green-1, green-7 and only red-1 digits are seen during training.

| Method | Red-7 Test Acc. (1) | Red-7 Test Acc. (2) |
| --- | --- | --- |
| ERM | 100% | 07% |
| GroupDRO | 100% | 26% |
| REx | 99% | 00% |
| CDANN + CenterLoss | 100% | **98%** |

**Supplementary Table 2:** Cross-tissue organ prediction on held-out patients.

| Method | Train Acc. | Test Acc. | Generalization Gap (Train-Test) |
| --- | --- | --- | --- |
| ERM | 0.978 | 0.682 | 0.296 |
| CDANN + Center Loss | 0.967 | **0.795** | 0.172 |

**Supplementary Table 3:**
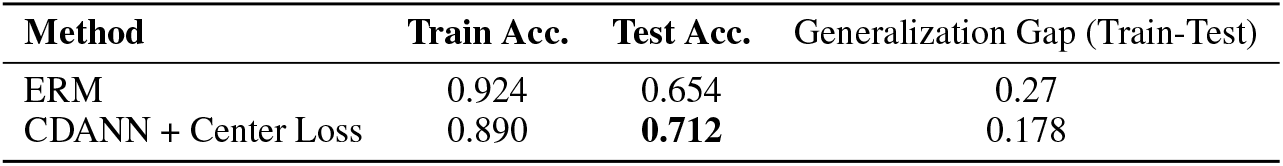
COVID-19 severity prediction on held-out cities.

| Method | Train Acc. | Test Acc. | Generalization Gap (Train-Test) |
| --- | --- | --- | --- |
| ERM | 0.924 | 0.654 | 0.27 |
| CDANN + Center Loss | 0.890 | **0.712** | 0.178 |

## 6.7 Supplementary Figures

**Supplementary Figure 1:**
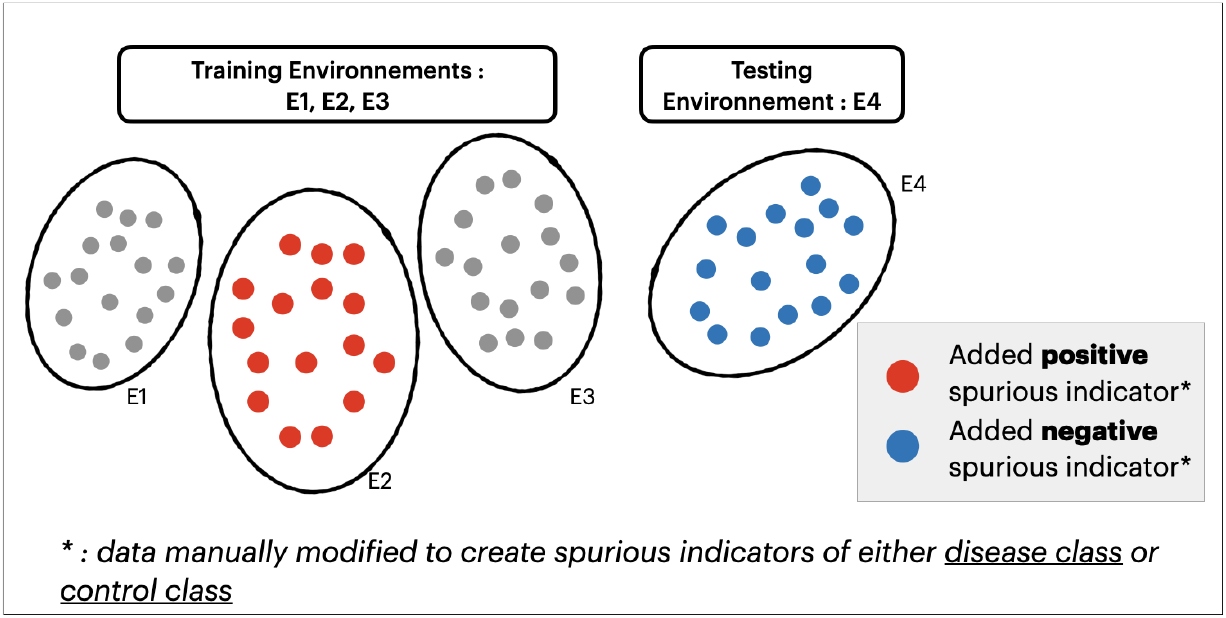
Schematic of environment setup. Spurious features may vary across environments: some are spurious-aligned (Env 2), neutral (Env 1-3), or spurious-inverted (Env 4). The goal is to generalize to held-out domains (Env 4) with different spurious associations.

**Supplementary Figure 2:**
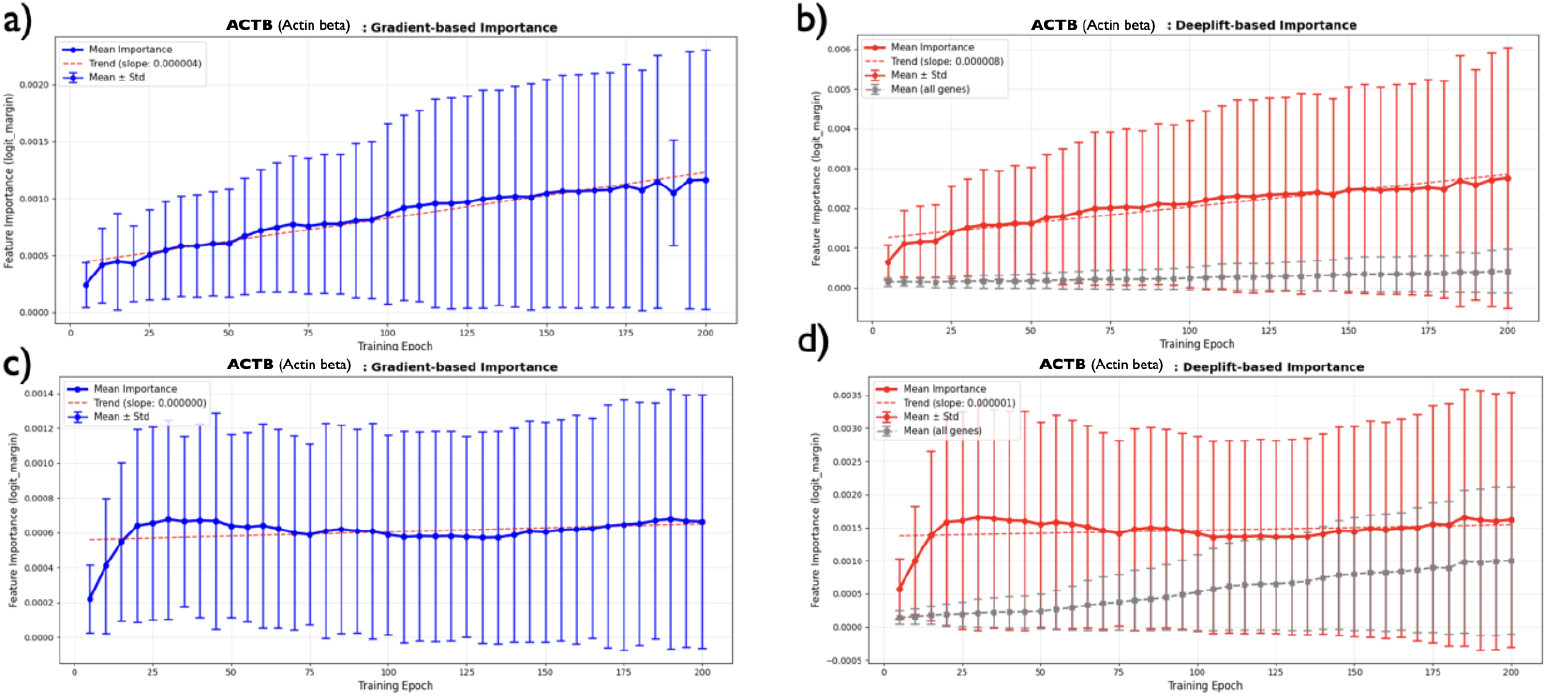
Attribution of a gene with synthetically-added spurious effects (*ACTB*) under different training strategies. **(a)** Gradient-based importance when the model is trained with ERM. **(b)** DeepLIFT-based importance when the model is trained with ERM. **(c)** Gradient-based importance when the model is trained with CDANN + CenterLoss. **(d)** DeepLIFT-based importance when the model is trained with CDANN + CenterLoss. Together, panels (a–b) show that under ERM the spurious gene gradually gains importance over epochs, indicating the model increasingly relies on the spuriously correlated signal. In contrast, panels (c–d) show that with CDANN + CenterLoss, the spurious gene remains near baseline attribution and is effectively disregarded, demonstrating improved robustness to spurious correlations.

**Supplementary Figure 3:**
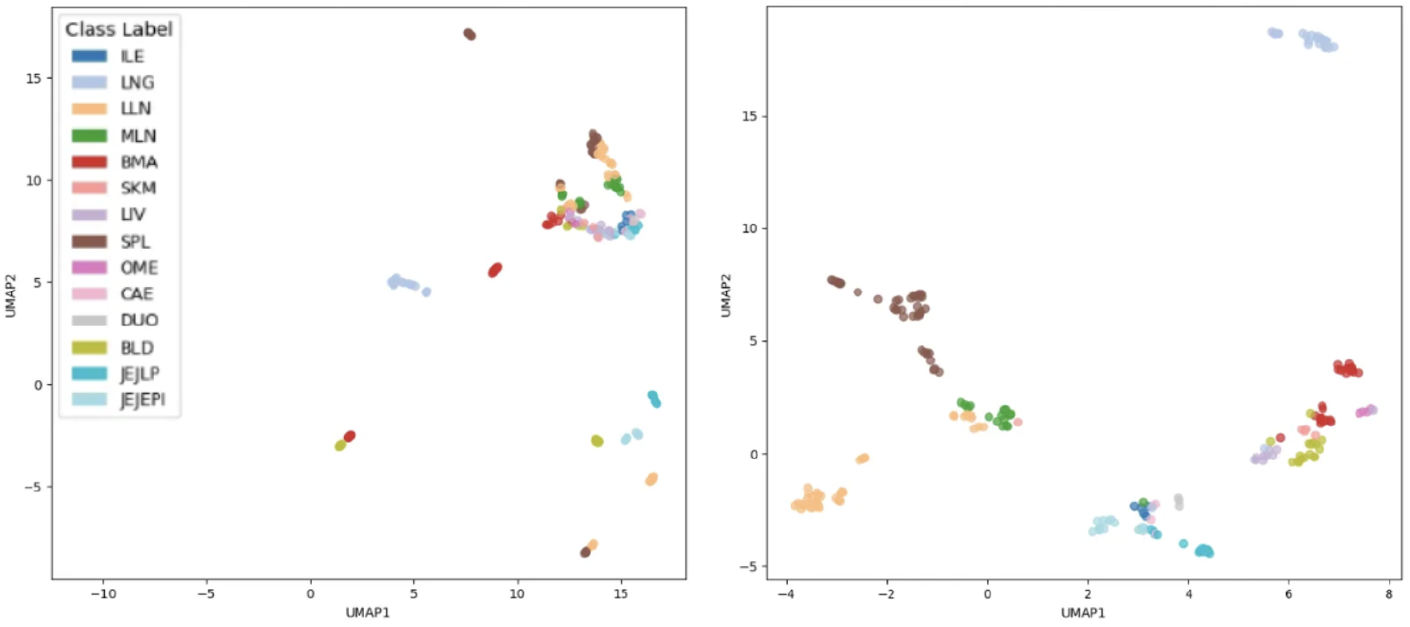
UMAP visualizations of patient embeddings colored by organ of origin. (left) Embeddings obtained from a model trained with ERM, showing limited separation between tissue types. (right) Embeddings from a model trained with CDANN + CenterLoss, displaying clearer clustering by tissue and improved disentanglement of donor-specific features.

**Supplementary Figure 4:**
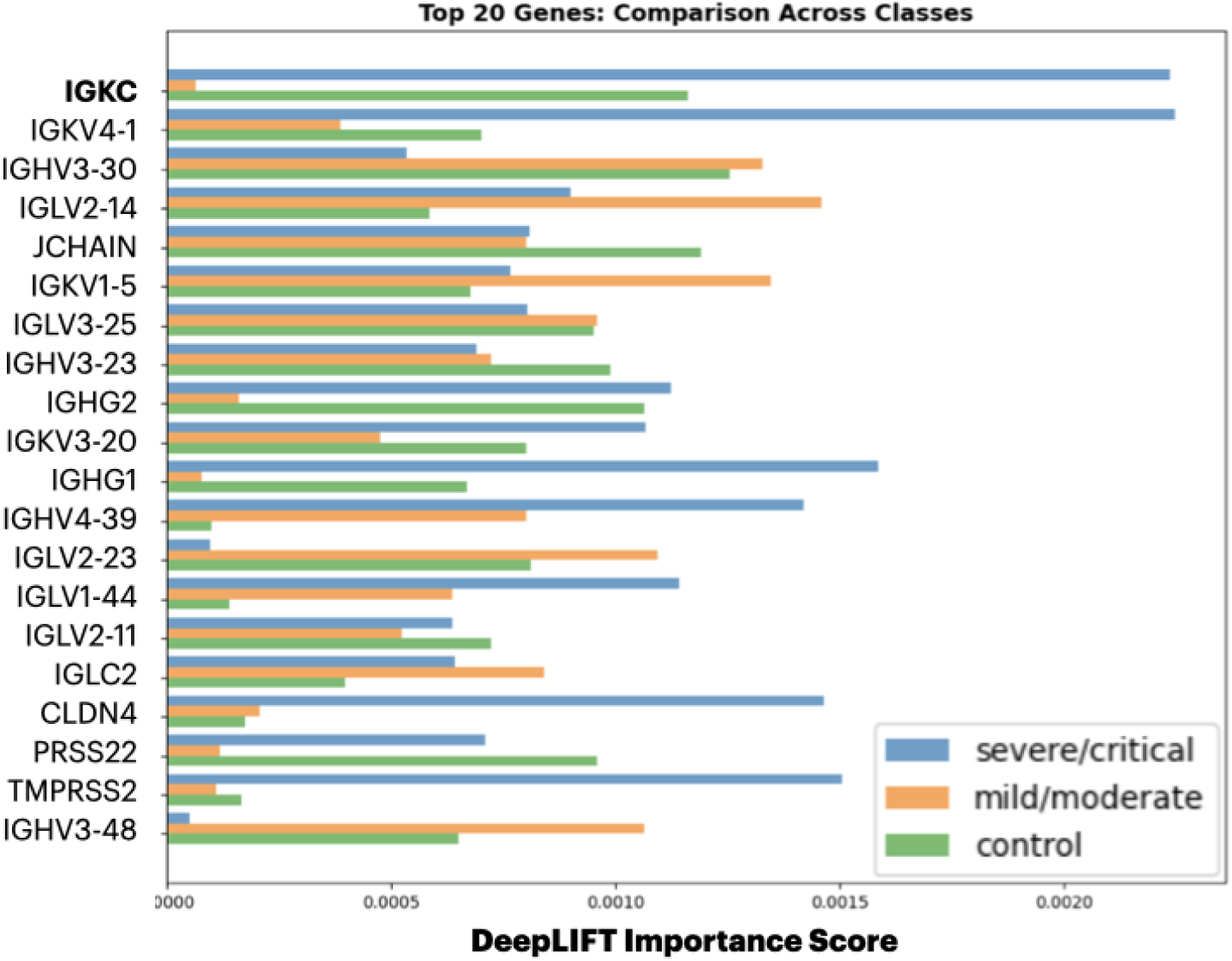
The top 20 genes by maximum DeepLIFT-based gene importance across disease severity classes for COVID-19 classification. The top gene is *IGKC*, also known as immunoglobulin *κ* constant.

